# The effects of multi-echo fMRI combination and rapid *T*_*2*_*\**-mapping on offline and real-time BOLD sensitivity

**DOI:** 10.1101/2020.12.08.416768

**Authors:** Stephan Heunis, Marcel Breeuwer, César Caballero-Gaudes, Lydia Hellrung, Willem Huijbers, Jacobus FA Jansen, Rolf Lamerichs, Svitlana Zinger, Albert P Aldenkamp

**Author notes:** Corresponding author: Stephan Heunis.

## Abstract

A variety of strategies are used to combine multi-echo functional magnetic resonance imaging (fMRI) data, yet recent literature lacks a systematic comparison of the available options. Here we compare six different approaches derived from multi-echo data and evaluate their influences on BOLD sensitivity for offline and in particular real-time use cases: a single-echo time series (based on Echo 2), the real-time *T*_*2*_*\**-mapped time series (*T*_*2*_*\*FIT*) and four combined time series (*T*_*2*_*\**-weighted, tSNR-weighted, TE-weighted, and a new combination scheme termed *T*_*2*_*\*FIT*-weighted). We compare the influences of these six multi-echo derived time series on BOLD sensitivity using a healthy participant dataset (N=28) with four task-based fMRI runs and two resting state runs. We show that the *T*_*2*_*\*FIT*-weighted combination yields the largest increase in temporal signal-to-noise ratio across task and resting state runs. We demonstrate additionally for all tasks that the *T*_*2*_*\*FIT* time series consistently yields the largest offline effect size measures and real-time region-of-interest based functional contrasts. These improvements show the possible utility of multi-echo fMRI for studies employing real-time paradigms, while caution is still advised due to decreased tSNR of the *T*_*2*_*\*FIT* time series. We recommend the use and continued exploration of *T*_*2*_*\*FIT* for offline task-based and real-time fMRI analysis. Supporting information includes: a data repository (https://dataverse.nl/dataverse/rt-me-fmri), an interactive web-based application to explore the data (https://rt-me-fmri.herokuapp.com/), and further materials and code for reproducibility (https://github.com/jsheunis/rt-me-fMRI).

## 1. Introduction

In functional magnetic resonance imaging (fMRI), *T*_*2*_*\**-weighted MRI sequences use the blood oxygen level-dependent (BOLD) signal as a proxy for neuronal activity. Our ability to infer accurate information about neuronal processes is influenced by the sensitivity with which we can capture these BOLD changes and subsequently delineate its sources of variance. Improved sensitivity is particularly important for real-time use cases, such as adaptive experimental paradigms, real-time quality control, or fMRI neurofeedback, where BOLD changes are quantified and used as they are acquired without the benefit of a full dataset or the requisite amount of post-processing time. It is well known that optimum sensitivity of single-echo fMRI is achieved at an echo time (TE) close to the apparent tissue *T*_*2*_*\**-value at baseline (Menon et al., 1993), which also underlies an inherent drawback of *T*_*2*_*\**-weighted sequences. Location-specific BOLD sensitivity is suboptimal since *T*_*2*_*\** varies across tissue types and brain regions (Peters et al., 2007), which can result in spatial variability in the detection of task-related activation patterns. Furthermore, magnetic susceptibility gradients on a macroscopic level result in image defects such as signal dropout and distortion, which is pronounced in the ventromedial prefrontal, orbitofrontal, the medial temporal and the inferior temporal lobes (Devlin et al., 2002). Additionally, the complex interplay of blood flow, blood volume and magnetic susceptibility effects can be influenced strongly by system- and participant-level noise sources, thus confounding the BOLD signal.

An advancement that has shown promise in making inroads into these drawbacks is multi-echo fMRI. Several studies have shown benefits of offline denoising based on multi-echo independent component analysis (MEICA; Kundu et al. 2012) for both resting state (e.g. Olafsson et al., 2015; Dipasquale et al., 2017) and task-based fMRI data (e.g., Lombardo et al., 2016; Gonzalez-Castillo et al., 2016; Moia et al., 2020). Echo combination via weighted summation is a critical step in multi-echo post-processing that has been reported to increase temporal signal-to-noise ratio, decrease signal drop-out, and improve activation extent for task-analysis (Poser et al., 2006). Posse et al. (1999) proposed several echo combination schemes, including simple echo summation (i.e. equal weights) and weighting echoes by their relative expected BOLD contrast contribution (i.e. *T*_*2*_*\**), which would require a numerical or fitted estimation of *T*_*2*_*\**. Other possible weighting schemes include optimised scalar weights, TE-weighted combination, and tSNR-weighted combination (also termed the PAID method) proposed by Poser et al. (2006). A theoretical framework for optimizing multi-echo combination has also been proposed by Gowland and Bowtell (2007). However, the relative benefits of all available combination schemes remain unclear.

With access to multiple data samples along the decay curve, multi-echo allows quantification of the effective transverse relaxation parameter *T*_*2*_*\** (decay time) or *R*_*2*_*\** (its inverse, decay rate), and *S*_*0*_ (initial net magnetization). This form of quantitative *T*_*2*_*\**-mapping (such as described by Weiskopf et al., 2013) acquires multiple closely spaced echoes followed by a data fitting procedure that yields a static, baseline *T*_*2*_*\**-map. In the context of functional imaging, however, temporal or per-volume *T*_*2*_*\**-mapping is also feasible, with the core benefit being the separation and quantification of *T*_*2*_*\** and *S*_*0*_ changes (from baseline) during stimulated neuronal activation. Such real-time use cases of multi-echo data have been reported, starting with Posse et al.’s (1998) single-shot, multi-echo spectroscopic imaging sequence that quantified region-specific *T*_*2*_*\** changes during olfactory and visual tasks, and which reported a larger functional contrast (up to 20% increase in the visual cortex) compared to standard EPI data. Several developments followed, including measuring single-event related brain activity (Posse et al., 2001), whole brain *T*_*2*_*\**-mapping at 1.5T using a linear combination of echoes (Hagberg et al., 2002), later with added gradient compensation (Posse et al., 2003), and a multi-echo EPI sequence at 3T with real-time distortion correction (Weiskopf et al., 2005). Rapid *T*_*2*_*\**-mapping has also been a useful tool in studying the interplay between cerebral blood flow, blood volume and blood oxygenation, particularly in combination with contrast agents (see, for example: Scheffler et al., 1999; Schulte et al., 2001; Pears et al., 2002). In real-time fMRI neurofeedback, some examples of multi-echo use are reported specifically for improving signal gains in regions such as the amygdala, including Posse et al. (2003) which uses *T*_*2*_*\**-weighted echo summation and Marxen et al. (2016), which uses scalar TE-dependent weights pre-selected to yield an average *T*_*2*_*\**-value of 30ms in the amygdala.

Although methodological studies have reported the benefits of multi-echo fMRI combination, a comprehensive evaluation of its practical benefits is lacking. Specifically, a variety of combination methods exist that can lead to both offline and real-time improvements in BOLD sensitivity, but there has been no systematic comparison between such methods. Additionally, per-volume *T*_*2*_*\**-mapping forms a necessary step in established multi-echo-based methods, but recent literature has not explored its value for task fMRI analysis. Consequently, this study has two main goals: (1) to explore the differences in BOLD sensitivity, both offline and per-volume, between time series of standard single echo EPI, per-volume estimated *T*_*2*_*\*FIT*, and multi-echo-combined time series (including tSNR-weighted, *T*_*2*_*\**, TE-weighted, and *T*_*2*_*\*FIT*-weighted); and (2) to explore the *T*_*2*_*\*FIT* time series as an alternative to single-echo or multi-echo-combined time series for offline and real-time fMRI analysis. We investigate these aims for whole brain data in separate task paradigms, eliciting responses to motor and emotion processing tasks and mental versions thereof, and during resting state. To quantify differences, we employ several metrics such as tSNR, task activity effect size, region-of-interest based temporal percentage signal change (tPSC), functional contrast, and temporal contrast-to-noise ratio (tCNR).

## 2. Multi-echo fMRI relaxation and combination

Multi-echo fMRI sequences acquire multiple whole brain functional images at discrete echo times (TE) after a single transverse excitation pulse of the scanner. This is done within the standard repetition time (TR) which then yields multiple echoes per volume. The relaxation of the fMRI signal in a given voxel after transverse excitation, assuming a mono-exponential decay model, is given as:

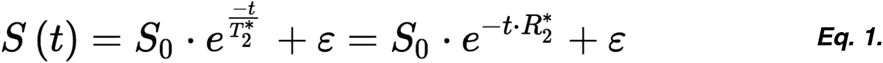

with *S(t)* being the time-decaying fMRI signal, *S*_*0*_ being the tissue magnetization directly after transverse excitation, and *T*_*2*_*\** being the local tissue transverse relaxation (i.e. decay time) constant (the inverse of the decay rate, *R*_*2*_*\**). Per-voxel estimates of *S*_*0*_ and *T*_*2*_*\** (depicted below in Fig. 1) can be derived using a log-linear regression estimation and the available echo times (*t*_*1*_ to *t*_*n*_, where *pinv* is the pseudo-inverse *log* the natural logarithm):

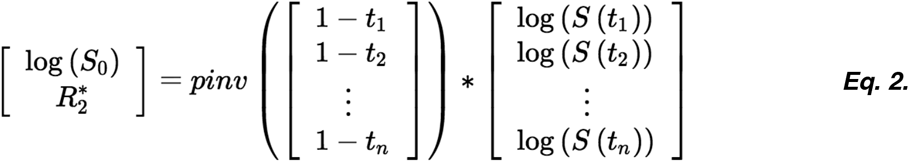

**Figure 1.**
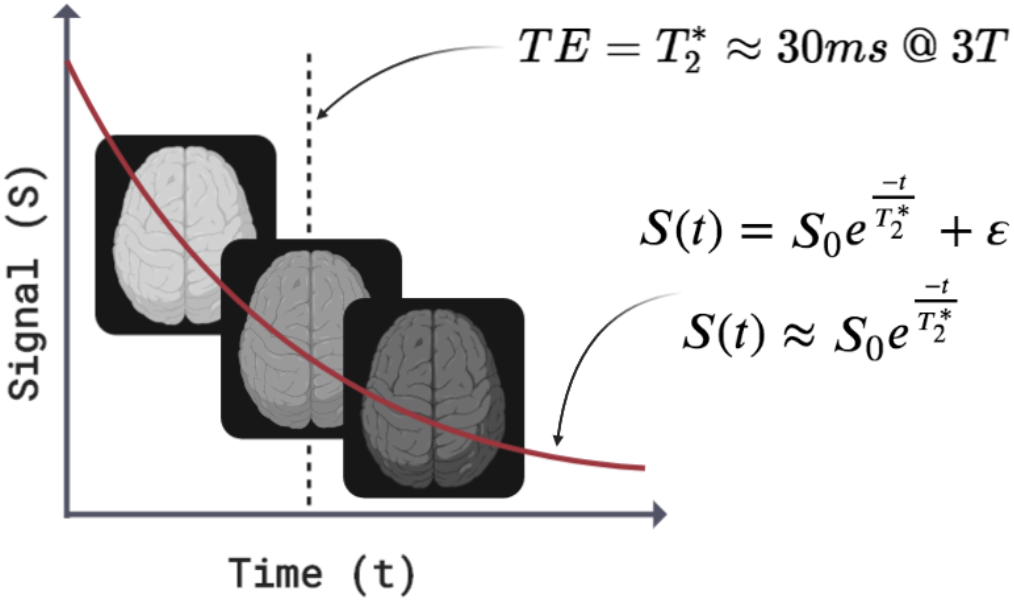
A representation of mono-exponential signal decay showing diminishing image intensity along three echoes. The second echo is sampled at the optimum echo time equal to average grey matter T_2_*, standard for single echo fMRI. The equation for the red, mono-exponential decay curve is provided (Eq. 1).

The mathematics of all widely used multi-echo combination schemes are based on the underlying concepts of data weighting, summation and averaging. In the supplementary online material, we provide a thorough background of these concepts along with explanatory equations S1 through S6. Importantly, the multi-echo combination schemes presented below use the convention of weighted summation with normalized weights. This implies that (1) all weights are normalized such that their sum equals 1, then (2) each normalized weight is multiplied by its corresponding data point, then (3) these products are summed to produce the weighted summation.

### Simple echo summation

Simple echo summation assumes equal weights for all echoes (totalling *N*), which is calculated for an individual echo *n* as:

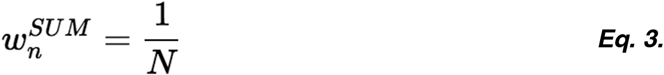

### T_2_*-weighted combination

The *T*_*2*_*\**-weighted combination scheme used by Posse et al. (1999) and termed “optimal combination” by Kundu et al. (2012), calculates the individual echo weights *w*_*n*_ per voxel as:

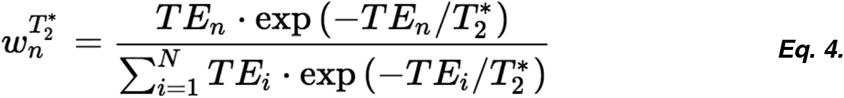

### tSNR-weighted combination

The PAID method put forward by Poser et al. (2006) uses the voxel-based tSNR measured at each echo (*tSNR*_*n*_) as the weights:

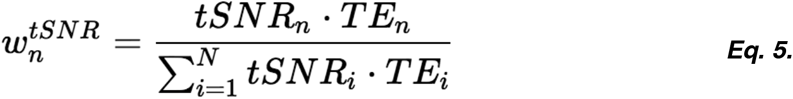

### TE-weighted combination

Purely using each echo’s echo time, *TE*_*n*_, as the weight for that echo has also been suggested (Posse et al, 1999). In this case, the same scalar value is used as the weighting factor for all voxels of a specific echo:

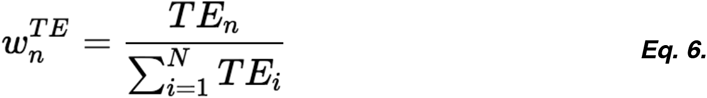

Similarly, a range of scalar values can be used as echo-dependent weighting factors, usually optimised according to study-specific criteria. For example, Marxen et al. (2016) selected scalar weights in order to yield an average *T*_*2*_*\** value of 30 ms in their region of interest (the amygdala). In such a case, the predefined scalar weights *{SW*_*1*_, *SW*_*2*_, …, *SW*_*N*_*}* can be normalized as:

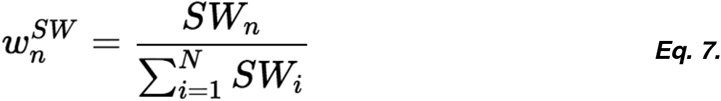

### T_2_*FIT-weighted combination

Finally, as proposed in the introduction, real-time *T*_*2*_*\**-mapping is made possible when using multi-echo fMRI. Here, the per-volume estimation of *T*_*2*_*\** at each voxel, termed *T*_*2*_*\*FIT(t)*, can also be used as the weighting factor in a per-volume echo combination scheme:

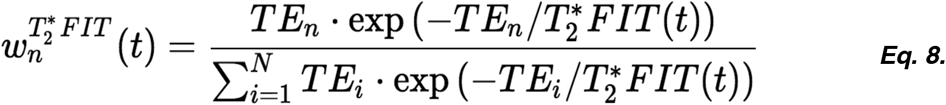

The per-volume nature of this echo combination scheme makes it ideal for use in both offline and real-time applications, when an priori *T*_*2*_*\**-map is not available or not preferred. To the best of our knowledge, this *T*_*2*_*\*FIT*-weighted combination approach has not been described previously in the literature

In the methods and results presented in this work, we compare metrics derived from standard single echo fMRI analysis to metrics derived from analysing *T*_*2*_*\**-weighted, tSNR-weighted, TE-weighted, *T*_*2*_*\*FIT*-weighted, and the *T*_*2*_*\*FIT* parameter time series, in both offline and per-volume scenarios.

## 3. Methods

In-depth descriptions of the participant details, ethics approval, experimental design, MRI protocol, preprocessing, and data quality can be accessed in the related data article (Heunis et al., 2020a). Summarising statements are provided below for the sake of completeness.

### 3.1. Participants

MRI and physiology data were collected from N=28 participants (male=20; female=8; age = 24.9 ± 4.6 mean + standard deviation). The study was approved by the local ethics review board and all participants gave written consent for their data to be collected, processed and shared in accordance with a GDPR-compliant procedure.

### 3.2. Experimental design

A total of seven MRI acquisitions were collected during a single scanning session per participant. These acquisitions include, in order of acquisition:

1. A T1-weighted anatomical scan
2. *rest_run-1*: the first resting state run, eyes fixated on a white cross
3. *fingerTapping*: a right hand finger tapping functional task
4. *emotionProcessing:* a matching-shapes-and-faces functional task
5. *rest_run-2*: the second resting state run, eyes fixated on a white cross
6. *fingerTappingImagined:* an imagined finger tapping functional task
7. *emotionProcessingImagined:* a functional task to recall an emotional memory

All four task paradigms followed an ON/OFF boxcar design, starting with the OFF condition, with both conditions lasting 10 volumes (= 20 s at TR = 2 s). The control (i.e. OFF) condition for the *fingerTapping* task was to focus on a small white cross on a black screen; for the *emotionProcessing* task the control condition was the shape-matching block; and for the *fingerTappingImagined* and *emotionProcessingImagined* tasks the control conditions were counting backwards, respectively, in multitudes of 7 and 9.

### 3.3. MRI protocol

MRI data were acquired on a 3 Tesla Philips Achieva scanner (software version 5.1.7) and using a Philips 32-channel head coil. A single T1-weighted anatomical image was acquired using a 3D gradient echo sequence (T1 TFE) with scanning parameters: TR = 8.2 ms; TE = 3.75 ms; flip angle = 8° field of view = 240×240×180 mm; resolution = 1×1×1 mm; total scan time = 6:02 min.

All six functional MRI scans were acquired using a multi-echo, echo-planar imaging sequence with scanning parameters: TR = 2000 ms; TE = 14,28,42 ms (3 echoes); number of volumes = 210 (excluding 5 dummy volumes discarded by the scanner); total scan time = 7:00 min (excluding 5 dummy volumes); flip angle = 90°; field of view = 224×224×119 mm; resolution = 3.5×3.5×3.5 mm; in-plane matrix size = 64×64; number of slices = 34; slice thickness = 3.5 mm; interslice gap = 0 mm; slice orientation = oblique; slice order/direction = sequential/ascending; phase-encoding direction = A/P; SENSE acceleration factor = 2.5. Parts of the cerebellum and brainstem were excluded for some participants to ensure full coverage of the cortex and subcortical areas of interest. Echo times, spatial resolution, and the SENSE factor were tuned with the aim of improving spatial resolution and coverage while limiting the TR to maximum 2000 ms, including a maximum number of echoes, and keeping the SENSE factor low to prevent SENSE artefacts.

In addition, cardiac and respiratory fluctuations were recorded during the functional scans, respectively using a pulse oximeter fixed to the participant’s left index finger, and a pressure-based breathing belt strapped around the participant’s upper abdomen. These were sampled at 500 Hz.

### 3.4. Data analysis

Data analysis consists of anatomical and functional preprocessing, definition and calculation of echo combination weights, multi-echo combination, time-series processing and calculation of comparison metrics. All analyses are done on an individual basis (i.e. participant-specific), unless otherwise stated, to describe the effects and facilitate the use of these methods in real-time fMRI use cases.

All processing steps below were done using the open source MATLAB-based and Octave-compatible *fMRwhy* toolbox (v0.0.1; *https://github.com/jsheunis/fMRwhy*), which has conditional dependencies:

- *SPM12* (r7771; https://github.com/spm/spm12/releases/tag/r7771; Friston et al., 2007)
- *bids-matlab (v*.*0*.*0*.*1, https://github.com/jsheunis/bids-matlab/releases/tag/fv0.0.1)*
- *Anatomy Toolbox* (v3.0; Eickhoff et al., 2005)
- *dicm2nii* (v0.2 from a forked repository; https://github.com/jsheunis/dicm2nii/releases/tag/v0.2)
- *TAPAS PhysIO* (v3.2.0; https://github.com/translationalneuromodeling/tapas/releases/tag/v3.2.0; Kasper et al., 2017)
- *Raincloud plots* (v1.1 https://github.com/RainCloudPlots/RainCloudPlots/releases/tag/v1.1; Allen et al., 2019).

All data analysis scripts can be accessed for reproducibility or reuse with attribution at https://github.com/jsheunis/rt-me-fMRI.

#### 3.4.1. Preprocessing

The basic anatomical and functional preprocessing pipeline applied to all data is described in detail in the data article, and in included:

1. Defining a functional template from Echo 2 of the first volume of the first resting state run.
2. Mapping prior data to the subject functional space, including:
  a. Coregistration of the anatomical image and atlas-based regions of interest to the functional template space, and resampling these to the functional resolution.
  b. Tissue-based segmentation the coregistered anatomical image and definition of binary maps for grey matter, white matter, cerebrospinal fluid (CSF) and the whole brain.
3. Basic functional preprocessing steps, including: estimating realignment parameters from the Echo 2 time series, running slice timing correction on all echo time series, applying realignment parameters to all echo time series, and applying spatial smoothing (7 mm isotropic, i.e. twice the voxel width) to all echo time series.
4. Generating data quality control metrics and visualizations to allow inspection of the quality of anatomical and functional data and their derivatives.

Two aspects of the preprocessing and analyses pipelines are worth highlighting in the context of this study. Firstly, while an important focus for this work is its application and utility in real-time scenarios, all processing was done offline, either on the full dataset or on a per-volume (i.e. simulated real-time) basis. This was viable since the study did not include any neurofeedback or real-time adaptive paradigms that would have required real-time computation and interaction. Secondly, the concept of “minimally processed” data was approached in accordance with suggestions from the tedana pipeline and its developers (see https://tedana.readthedocs.io/en/latest/index.html; Dupre et al., 2020), which states that minimal steps (including slice timing correction and 3D volume) should be applied to multi-echo data before decay parameter estimation or multi-echo combination.

#### 3.4.2. Data quality control

The *fMRwhy* toolbox has a BIDS-compatible data quality pipeline for functional and anatomical MRI, *fmrwhy_bids_workflowQC*, that can be run automatically for a full BIDS-compliant dataset. After running minimal preprocessing steps it generates a subject-specific HTML-report with quality control metrics and visualizations to allow inspection of the data and its derivatives. Individual reports can be accessed in the derivatives directory of the shared BIDS-compliant dataset of this study (see Heunis et al., 2020a for details). Additionally, a web-application named ***rt-me-fMRI*** is provided along with this work and accessible at: https://rt-me-fmri.herokuapp.com/. It can be used interactively to explore various summaries of data quality metrics, including distributions of framewise displacement (FD) and tSNR, and physiology recordings, as well as the results of this study.

None of the participant datasets were excluded after inspection of the included quality metrics, even in cases of more than average or severe motion (specifically sub-010, sub-020, and sub-021), since such data could still be useful for data quality related insights or for future denoising methods validation.

#### 3.4.3. Multi-echo combination

Existing weighting parameters or parameter maps are required to allow both offline and per-volume combination of multi-echo data. Of the previously reported options for weighting schemes given in Section 2.2, the simplest option used in this study is the echo time (Eq. 6) derived from the functional MRI protocol, which yields a TE-weighted combination. Other prior weighting parameters are calculated using the first resting state functional scan. For each minimally preprocessed echo time series of the resting state run, the time series mean and standard deviation are calculated. The mean divided by the standard deviation yields the temporal signal-to-noise ratio (tSNR), per echo, that is used as another weighting parameter (Eq. 5) described as the PAID method by Poser et al. (2006). Additionally, the mean images from the three echo time series are used to derive the per-voxel estimates of *S*_*0*_ and *T*_*2*_*\** assuming a mono-exponential decay model and using a log-linear regression estimation (Eq. 2). This baseline *T*_*2*_*\** map can be used for *T*_*2*_*\**-weighted combination (Eq. 4), described as optimal combination by Kundu et al. (2012). Lastly, the same log-linear regression that is applied to the time series mean images can also be applied to a single volume of any multi-echo data. This implies that the three echo images of any volume can be used as data points to estimate per-volume and per-voxel parameter maps, *S*_*0*_*FIT(t)* and *T*_*2*_*\*FIT(t)*, which in turn can be used for per-volume multi-echo combination (Eq. 8), hereinafter referred to as *T*_*2*_*\*FIT*-combination.

Multi-echo combination schemes are applied to all functional data excluding the first resting state run, from which prior baseline weight maps are derived. In sum, six time series are computed per functional run (as described in Fig. 2): Echo 2, *T*_*2*_*\**-combined, tSNR-combined, TE-combined, *T*_*2*_*\*FIT*-combined, and the *T*_*2*_*\**FIT time series.

**Figure 2:**
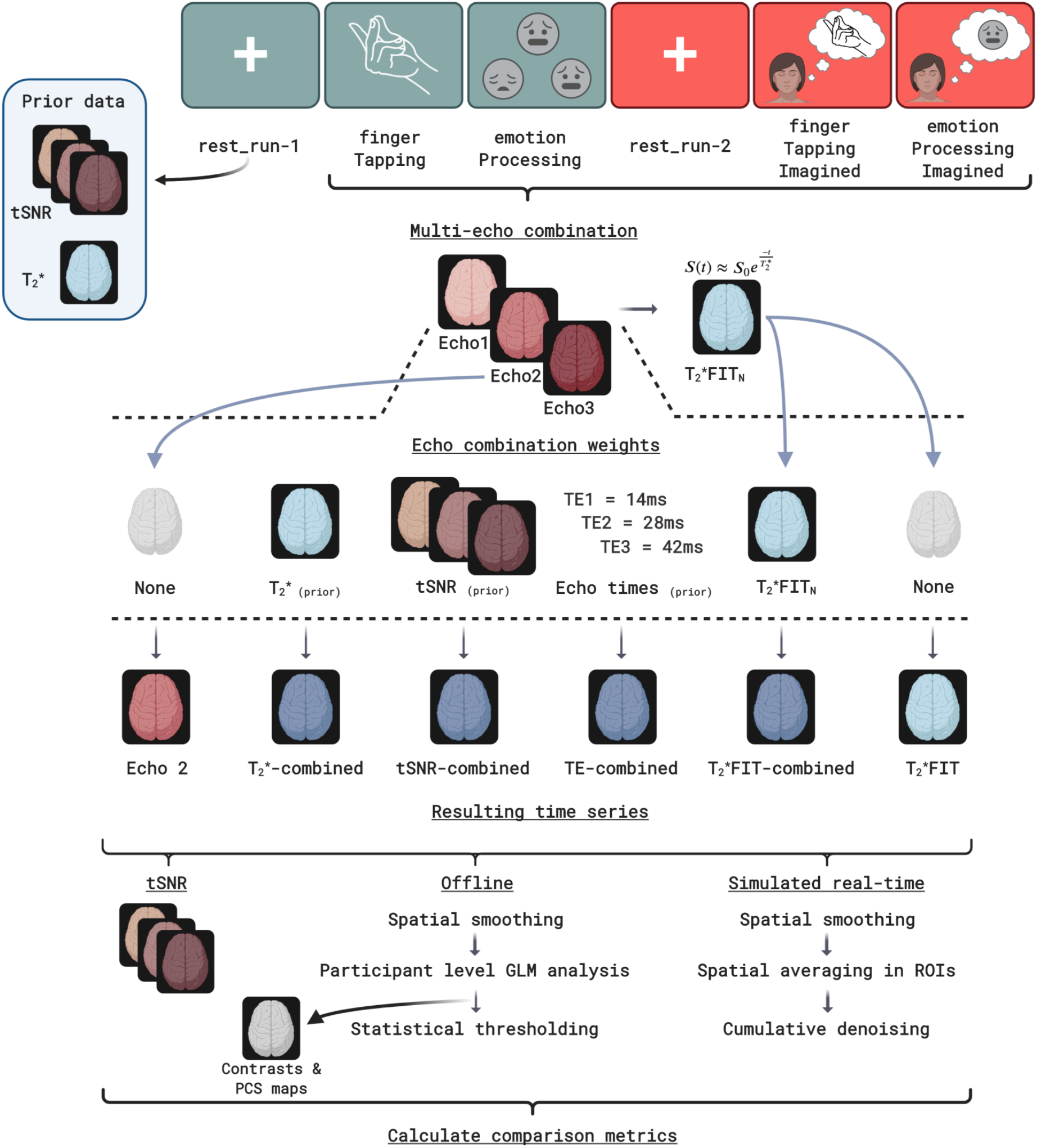
The analysis pipeline applied to the rt-me-fMRI dataset. Prior tSNR and T_2_* maps are derived from the first resting state run. For all other functional runs (top row), six steps are executed per volume after minimal preprocessing in order to yield resulting multi-echo-derived time series for comparison: (1) the 2nd echo time series is extracted without processing, (2) the baseline T_2_*-weighted combination, (3) the prior tSNR-weighted combination, (4) the TE-weighted combination, (5) the T_2_*FIT-weighted combination, and (6) T_2_*FIT time series. Following this, each of the six time series then undergoes offline and simulated real-time processing pipelines. The offline pipeline includes (in order): tSNR calculation, spatial smoothing, participant-level task analysis, calculation of percentage signal change effect sizes, and statistical thresholding of the participant-level contrast maps. The simulated real-time pipeline is run per volume for each time series and includes (in order): spatial smoothing, spatial averaging of the appropriate region-of-interest signals, and cumulative denoising (including detrending using linear and quadratic regressors).

#### 3.4.4. Time series processing

After computing the six time series per functional run (excluding the first resting state run), each resulting time series is processed as summarised in the bottom row of Fig. 2.

First, the tSNR of each time series is calculated prior to any further processing. Then, each time series is spatially smoothed using a Gaussian kernel with FWHM at 7 mm (i.e. double the voxel size). This is followed by participant-level GLM-based analysis of the four task runs. Task regressors included the main “ON” blocks for the *fingerTapping, fingerTappingImagined* and *emotionProcessingImagined* tasks, and both the separate “SHAPES” and “FACES” trials for the *emotionProcessing* task. Regressors not-of-interest for all runs included six realignment parameter time series and their derivatives, the CSF compartment time series, and RETROICOR regressors (both cardiac and respiratory to the 2nd order, excluding interaction regressors). Additional steps executed by SPM12 before beta parameter estimation include high-pass filtering using a cosine basis set and AR(1) autoregressive filtering of the data and GLM design matrix. Contrasts are applied to the task-related beta maps for the *fingerTapping, fingerTappingImagined* and *emotionProcessingImagined* tasks, and to the FACES, SHAPES, and FACES>SHAPES beta maps for the *emotionProcessing* task. In order to yield a standard measure of effect size, the parameter estimates or contrast maps are then used to calculate percentage signal change (PSC) using the method described by Pernet (2014) and given by:

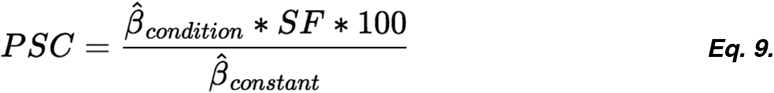

where *β*_*condition*_ and *β*_*constant*_ are parameter estimates corresponding to the relevant GLM regressors that are scaled with regards to the actual BOLD magnitude. To account for this the scaling factor, *SF*, is determined as the maximum value of a reference trial taken at the resolution of the super-sampled design matrix *X*_*ss*_ (where supersampling is typically done before convolution with the hemodynamic response function):

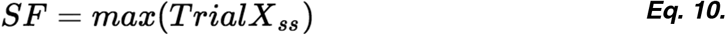

Statistical thresholding was applied to identify task-related clusters of activity by controlling the familywise error rate (FWE), with p < 0.05, and a voxel extent threshold of 0.

#### 3.4.5. Real-time analysis

Minimally processed time series are also analysed per-volume (using data acquired up to each volume in time) in order to explore multi-echo related BOLD sensitivity changes for real-time applications. Real-time analysis typically involves minimal processing (including 3D realignment), spatially averaging the signal within given ROIs, and additional per-volume denoising steps on the averaged signal. Here, we run a per-volume denoising process adapted from OpenNFT (Koush et al., 2017) on all task time series. This process is depicted in the bottom row of Fig. 2 and includes, in order: 1) Spatial smoothing using a Gaussian kernel with FWHM at 7 mm, 2) Spatial averaging of voxel signals with defined ROIs, and 3) Cumulative GLM-based detrending of the ROI signals, including linear and quadratic trend regressors. This then yields per-volume minimally denoised ROI-signals from which percentage signal change or another calculation can be used as the basis for the neurofeedback or real-time ROI signal.

#### 3.4.6. Comparison metrics

To explore the differences between various multi-echo combinations and standard single echo data, and to investigate the usefulness of the former over the latter, we employ several comparison metrics:

- **Temporal signal-to-noise ratio (tSNR)**, calculated as the voxel-wise time series mean divided by voxel-wise time series standard deviation. tSNR is an indicator of the amount of signal available from which to extract potentially useful BOLD fluctuations. Additionally, tSNR maps can be a robust visual indicator of increases or decreases in signal dropout.
- **Percentage signal change (PSC)**, of task-based contrast maps resulting from participant-level GLM analysis. PSC represents a standardised measure of effect size (which beta or contrast values are not) and is an indicator of the BOLD sensitivity of the data based on GLM analysis.
- **T-statistic values**, related to the task based contrast maps resulting from participant-level GLM analysis.
- **Temporal percentage signal change (tPSC)**, of the single echo, combined-echo and derived time series data of the task runs. This is calculated per voxel on minimally processed task data as the per-volume signal’s percentage signal change from the time series mean (or, for real-time scenarios, from the mean of the preceding baseline “OFF” block or the cumulative mean). These are then spatially averaged within the regions listed below to yield ROI-based time series. These time courses are similar to what would be calculated in real-time as the ROI-based neurofeedback signal, and their amplitudes can be an indicator of BOLD sensitivity.
- **Carpet plots for the whole brain and within regions of interest**. These plots show the voxel intensity over time in terms of tPSC and can be a valuable visual indicator of quality issues (Power, 2017) and also of amplitude differences (i.e. functional contrast) in ROIs.
- **Functional contrast:** of the ROI-based tPSC signals. To calculate the functional contrast in ROIs, the average tPSC in volumes classified as being part of “OFF” condition blocks are subtracted from the average signal in volumes classified as being part of each “ON” condition block. Visually, this corresponds to the average amplitude difference between conditions in the tPSC signal. The functional contrast is an indicator of the BOLD sensitivity of a signal based on both minimally processed and denoised data.
- **Temporal contrast-to-noise ratio (tCNR)** of the single echo, combined-echo and derived time series data of the task runs. To calculate the tCNR, the functional contrast in an ROI is divided by the time series standard deviation of the tPSC signal in the same ROI. This is related to both the tSNR and BOLD sensitivity. Where tPSC consists of time courses, tCNR provides a single summary value per voxel or region.

Extracting and spatially averaging voxel time series from specific regions is a common approach to exploring patterns of task-based activity in fMRI (Poldrack, 2006). This can be done both offline on a full dataset, and in real-time on the data as they are acquired. In this work, we explore and compare the above-mentioned metrics on both whole-brain and region-specific levels. Regions include:

- Grey matter (**GM**), white matter (**WM**) and cerebrospinal fluid (**CSF**) compartments. This allows quantifying, for example, whether combined multi-echo data changes a given metric similarly or differently across tissue types.
- Binary task-based clusters of voxels surviving conservative statistical thresholding (**FWE**). These clusters vary spatially per time series of a given task run and they represent the clusters of functionally most responsive voxels based on the underlying data but assuming a shared criteria (i.e. statistical threshold).
- A binary cluster of voxels that is a logical OR of the conservatively thresholded task based clusters of all six time series of a given task run (**FWE-OR**). This allows the comparison of metrics in a region that includes the voxels that are judged to be significantly active in *any time series*, thus removing time series-specific spatial bias.
- Atlas-based anatomical regions of interest (**Atlas-based ROI**) that have been mapped to individual anatomical scans and coregistered and resampled to the individual functional space. This allows quantification of the above metrics within an a priori defined ROI, thus excluding spatial bias introduced by statistical thresholding. The Atlas-based ROIs include the **left motor cortex** (for right-hand finger tapping), and the **bilateral amygdala** (for emotion processing).

## 4. Results

A web-application named ***rt-me-fMRI*** is provided alongside this work and accessible at: https://rt-me-fmri.herokuapp.com/. This browser-based application can be used interactively to explore the summary and participant-specific results presented below, and is intended to serve as supplementary material to this work.

### 4.1. Multi-echo decay

To illustrate signal decay and dropout as a function of echo time, a simple plot of the inferior slices of a single subject is given In Fig. 3. Signal decay can be seen clearly as the signal intensity diminishes from Echo 1 to Echo 3 (top to bottom) in all displayed slices. Signal dropout from Echo 1 through Echo 3 is particularly evident in the areas of the orbitofrontal and ventromedial prefrontal cortices, and the inferior and anterior temporal lobe (magenta arrows; slices 8, 10, 12) and the cerebellum (light blue arrows; slices 2, 4, 6).

**Figure 3:**
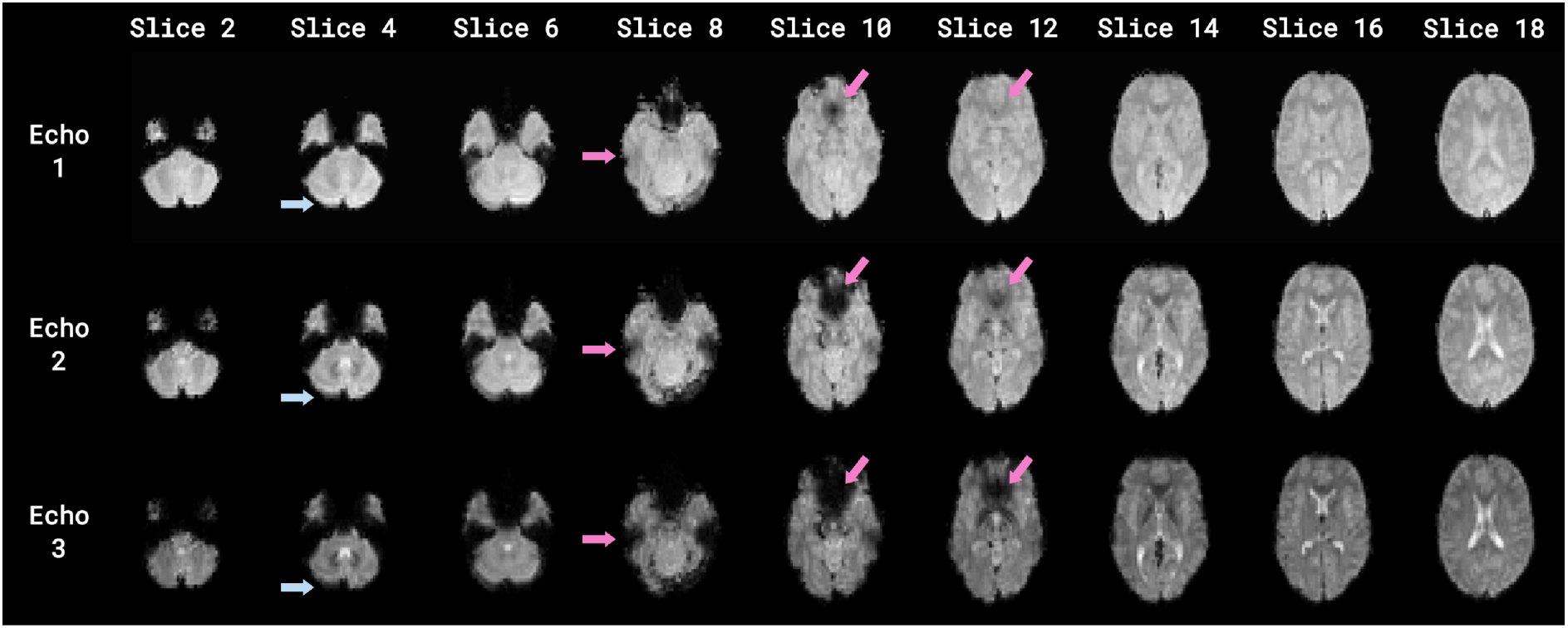
Signal decay along the three echoes (top to bottom) of a single volume. Signal decay is displayed across a selection of slices (horizontal axis). Signal dropout is clearly evident in the orbitofrontal and ventromedial prefrontal cortices and inferior and anterior temporal lobes (magenta arrows; slices 8, 10, 12) and the cerebellum (light blue arrows; slices 2, 4, 6).

### 4.2. Signal intensity, dropout, and temporal signal-to-noise ratio

We can visually inspect the effect on signal intensity and dropout when combining multi-echo data or deriving time series from it. Fig. 4A shows the mean of each of the six time series: Echo 2, *T*_*2*_*\**-weighted combination, tSNR-weighted combination, TE-weighted combination, *T*_*2*_*\*FIT*-weighted combination, and *T*_*2*_*\*FIT*.

**Figure 4:**
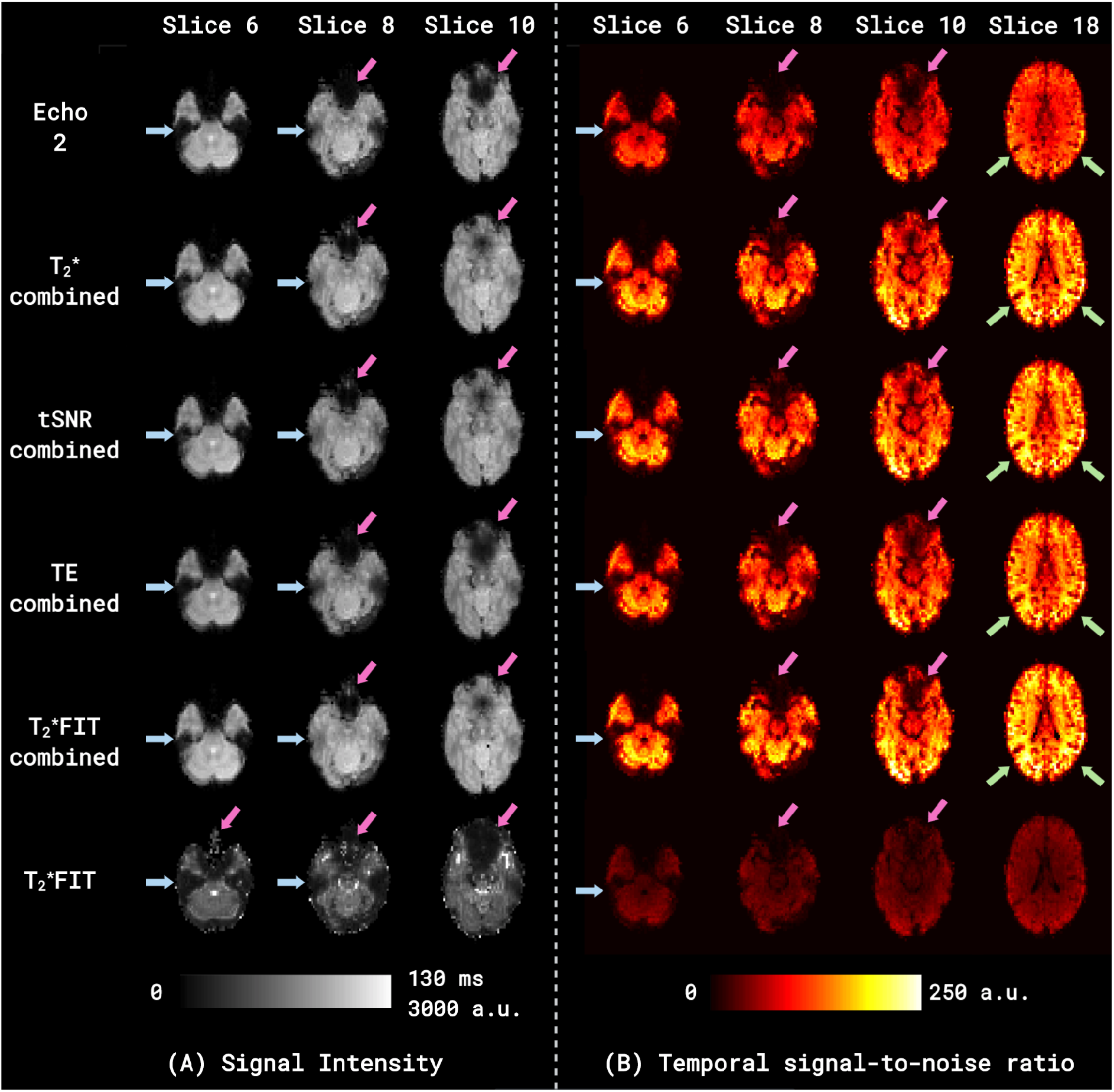
Signal intensity (Fig. 4A) and Temporal signal-to-noise ratio (Fig. 4B) shown in mean images for the time series in rows from top to bottom: Echo 2, T2*-weighted combination, tSNR-weighted combination, TE-weighted combination, T2*FIT-weighted combination, and T2*FIT. Scaling for Fig. 4A is given both for the T2*FIT signal (0-130 ms) and for all the other signals (0-3000 a.u.). All echo combination schemes, with the exception of TE-weighted combination, recover some signal lost due to dropout in the orbitofrontal and ventromedial prefrontal regions (magenta arrows; slices 8, 10) and inferior and anterior temporal regions (light blue arrows; slices 6, 8). Slight signal recovery in T2*FIT is visible in the orbitofrontal and ventromedial prefrontal regions (slice 6) although signal loss is more evident in slice 10. In Fig. 4B, all time series apart from T2*FIT show increases in tSNR (from Echo 2) in areas close to the bilateral temporal-occipital junction and towards the occipital lobe (green arrows; slice 18), which is more pronounced in the T2*FIT-weighted compared to the T2*-weighted and tSNR-weighted combinations, and less so in the TE-weighted combination.

It is evident that most echo combination schemes, with the exception of TE-weighted combination, recover some signal lost due to dropout in the orbitofrontal and ventromedial prefrontal regions (magenta arrows; slices 8, 10) and inferior and anterior temporal regions (light blue arrows; slices 6, 8). This signal recovery is further demonstrated in the tSNR maps provided in Fig. 4B, particularly by the magenta arrows showing areas of signal dropout in Echo 2 and subsequent recovery in combined and derived time series tSNR maps. Even the *T*_*2*_*\*FIT*, for which the tSNR is evidently much lower than all other time series including Echo 2, recovers some of the signal that is lost due to low BOLD sensitivity in the affected areas, although signal loss is also more evident (slice 10). Additionally, tSNR in areas close to the bilateral temporal-occipital junction and towards the occipital lobe (Fig. 4B, green arrows) appears to increase substantially for all combined time series vs. Echo 2. This is more pronounced in the *T*_*2*_*\*FIT*-weighted compared to the *T*_*2*_*\**-weighted and tSNR-weighted combinations, and less so in the TE-weighted combination.

To provide a more quantified view than these visualisations of signal intensity (Fig. 4A) and tSNR (Fig. 4B), distribution plots were created for grey matter tSNR values, both for subjects individually and for the whole group. Fig. 5 shows these distributions together as a ridge plot for *sub-001_task-rest_run-2*, from which we can see the mean tSNR increase for all combined time series compared to Echo 2, with the *T*_*2*_*\*FIT*-weighted combination showing the largest increase (36.14%) and the *T*_*2*_*\*FIT* time series showing a substantial decrease (−55.89%).

**Figure 5:**
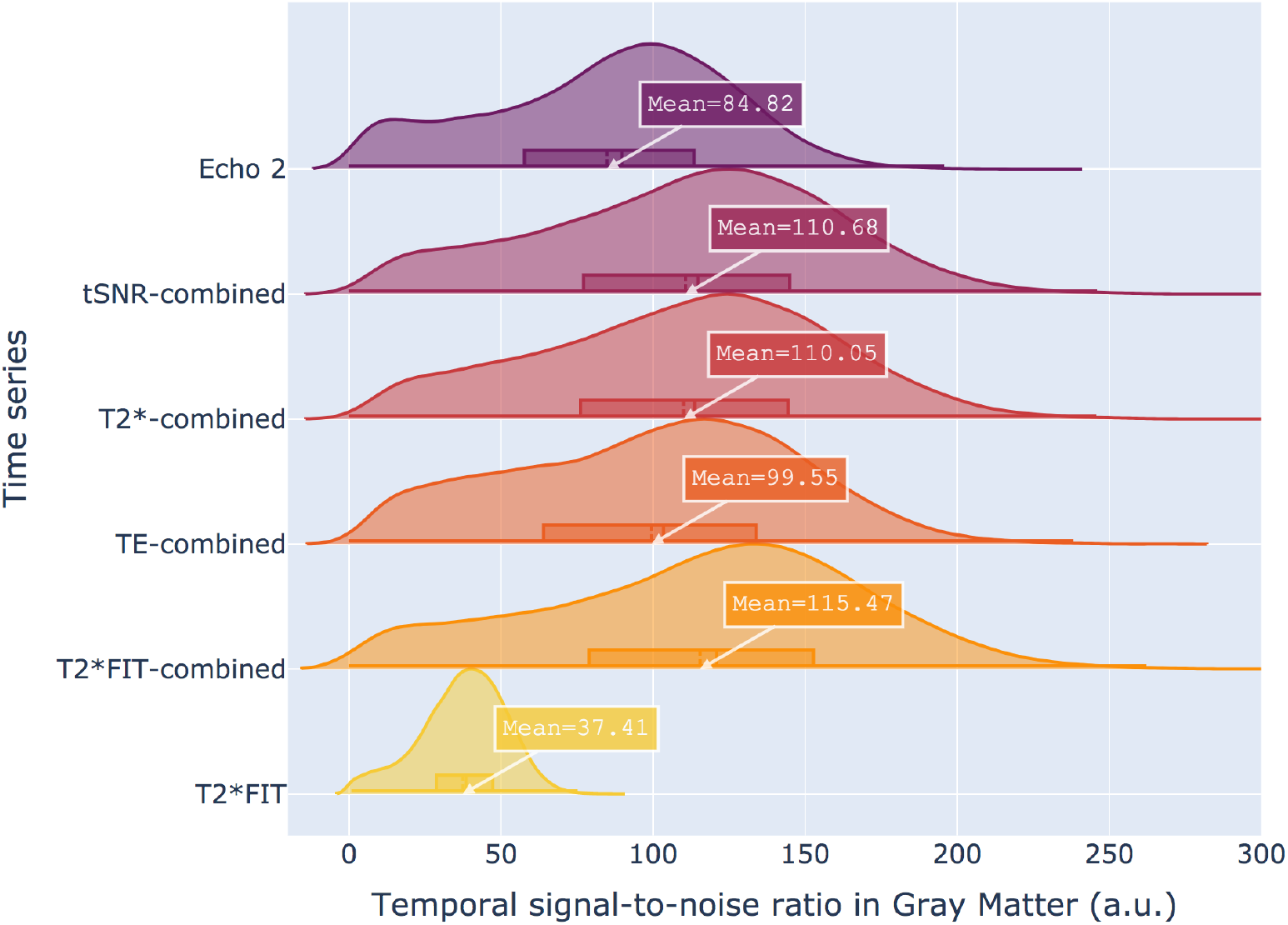
Distribution / ridge plots of whole brain grey matter temporal signal-to-noise ratio (tSNR) for a single run of a single subject. Plots are shown for the six time series in rows from top to bottom: Echo 2, tSNR-weighted combination, T2*-weighted combination, TE-weighted combination, T2*FIT-weighted combination, and T2*FIT. While the mean T2*FIT tSNR decreases from Echo 2, the tSNR of all other time series increase, with the T2*FIT-weighted combination showing the largest increase from 84.82 to 115.47 (36.14%).

The mean grey matter tSNR values are summarised on a group level for all subjects and all functional runs in Fig. 6. Fig. 6A displays mean grey matter tSNR for the whole brain, while Figs. 6B and 6C show the same for the left motor cortex and the bilateral amygdala, respectively. Fig. 6A shows the same relationship between tSNR values for single echo, combined echo, and derived time series as was seen for a single run of the single subject in Fig. 5, i.e. a mean tSNR increase for all combined time series compared to Echo 2, with the *T*_*2*_*\*FIT*-weighted combination showing the largest increase (a comparable 36.95%).

**Figure 6:**
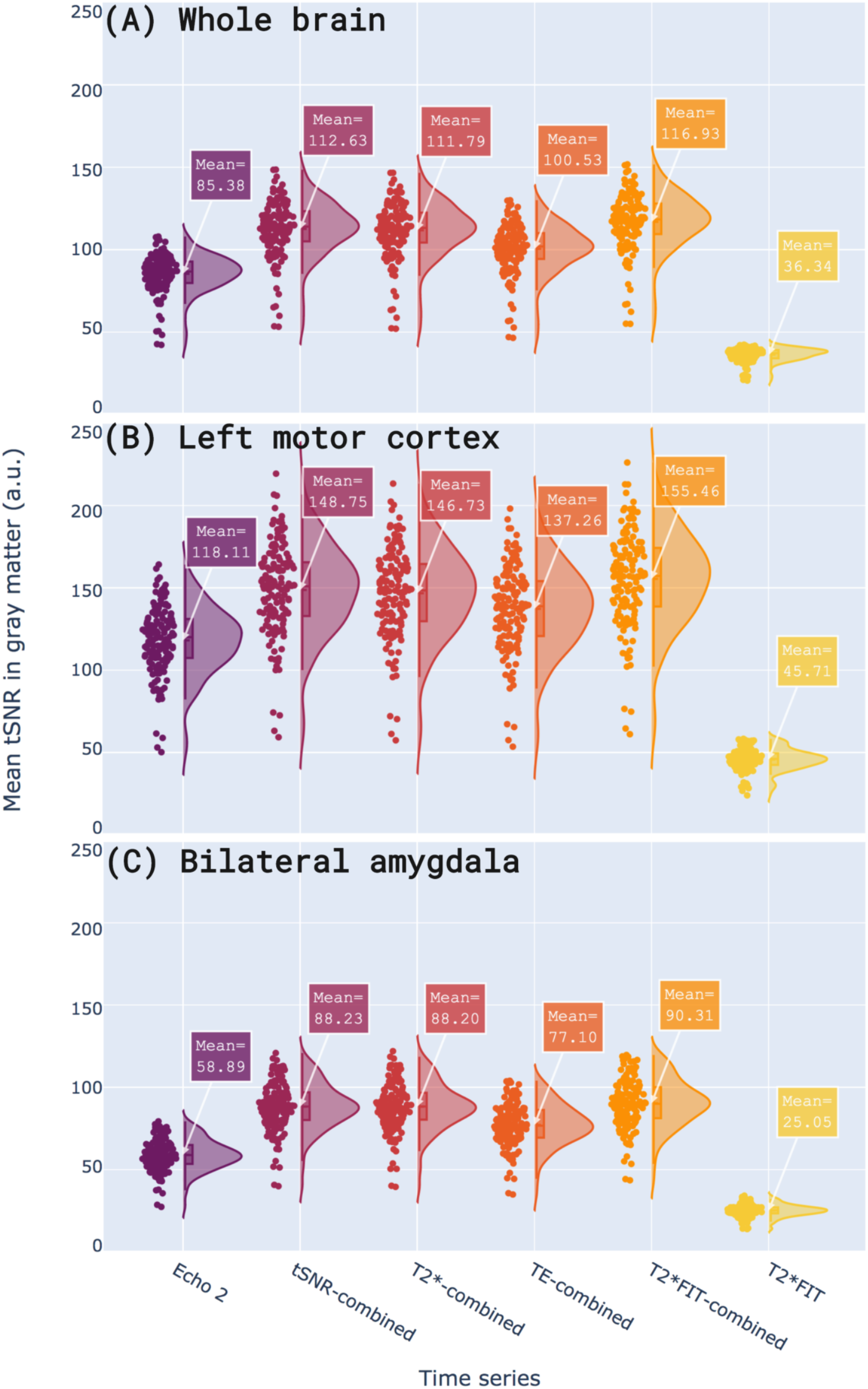
Distribution / ridge plots of mean grey matter temporal signal-to-noise ratio (tSNR) over all participants and all runs. Plots are shown for (A) the whole brain, (B) the left motor cortex, and (C) the bilateral amygdala, each displaying a distribution for the six time series from left to right: Echo 2, tSNR-weighted combination, T2*-weighted combination, TE-weighted combination, T2*FIT-weighted combination, and T2*FIT. In all regions, the mean T2*FIT tSNR decreases from Echo 2 while the tSNR of all other time series increase, with the T2*FIT-weighted combination showing the largest increase in all regions. Notably, tSNR increases for all the combined echo time series are more substantial in the amygdala (C) than the other regions (A, B).

This increase in the tSNR of *T*_*2*_*\*FIT*-weighted combination replicates results that we previously reported on a different dataset (Heunis et al., 2019). This relationship also repeats for different regions, as can be seen for the left motor cortex (Fig. 6B) and the bilateral amygdala (Fig. 6C).

Note, however, that the mean tSNR values increase differentially based on the region. For the *T*_*2*_*\*FIT*-weighted combination, for example, whole brain data show a mean tSNR increase of 36.95%; the left motor cortex shows a mean tSNR increase of 31.63%; and the bilateral amygdala shows a mean tSNR increase of 53.35%. Other combined time series show percentage increases following the same pattern, i.e. a larger tSNR increase for the amygdala than for the whole brain or motor cortex. This could be explained by the baseline *T*_*2*_*\*-*values in the motor cortex and the whole brain being closer to the time Echo 2 (28 ms) than the *T*_*2*_*\*-* values in the amygdala, i.e. that the *T*_*2*_*\*-*weighting of Echo 2 in those regions is already closer to optimal than the weighting of Echo 2 in the amygdala. This suggests that the amygdala and similarly affected areas with *T*_*2*_*\*-*values that are different from the average have more to gain from the multi-echo combination process.

Another noteworthy aspect is the low signal intensity and low tSNR of the *T*_*2*_*\*FIT* time series. The low signal intensity is explained by the fact that *T*_*2*_*\*FIT* values correspond to quantified units (ms) that are expected to be in a certain range (∼ 0 to 120 ms for the human brain at 3T, Peters et al., 2007), while the intensity of the standard single and combined echo images are in analogue units determined by MRI hardware and software. The low tSNR of the *T*_*2*_*\*FIT* time series could be explained by an increase in time series standard deviation resulting from the log-linear fitting procedure on noisy data and only using the three echoes to fit the mono-exponential decay model per volume. This increase in time series noise becomes evident below when investigating temporal percentage signal change.

### 4.4. Effect sizes and T-statistics

Figure 7 shows distribution plots (over all subjects) of the mean PSC values within the respective **FWE-OR** clusters for all task runs: *fingerTapping* (Fig. 7A), *fingerTappingImagined* (Fig. 7B), *emotionProcessing* (Fig. 7C), and *emotionProcessingImagined* (Fig. 7D). It is evident from Fig. 7A through 7D that the effect sizes show a substantial increase for the *T*_*2*_*\*FIT* time series (from Echo 2) in all tasks (respectively 87.91%, 67.86%, 13.51%, and 43.28%), while displaying a similar or decreased mean effect size for all combined times series. Data in the supplementary browser-based application also shows that this increase for *T*_*2*_*\*FIT* is more pronounced when looking at the effect sizes within their respective **FWE** clusters (i.e. different activated voxels for each multi-echo derived time series, although mostly overlapping), which one should be wary of overinterpreting given the inherent circularity of re-analysing data in voxels that previously passed a significance threshold using the same data. On the other hand, this result is less pronounced for the time series effect sizes within their respective **atlas-based** regions of interest, mainly resulting in a longer tailed distribution of mean PSC values for the *T*_*2*_*\*FIT* time series. In some participants the mean PSC values of the *T*_*2*_*\*FIT* time series even show a slight decrease. These decreases in PSC disappear when looking at peak effect sizes, as opposed to mean effect sizes, in the respective regions (**FWE, FWE-OR**, and **atlas-based**). Further differences in effect size increases/decreases, especially between the Echo 2 time series and the *T*_*2*_*\*FIT* time series, and within the **FWE** versus the **atlas-based** regions of interest, can be inspected in depth using the supplementary browser-based application.

**Figure 7:**
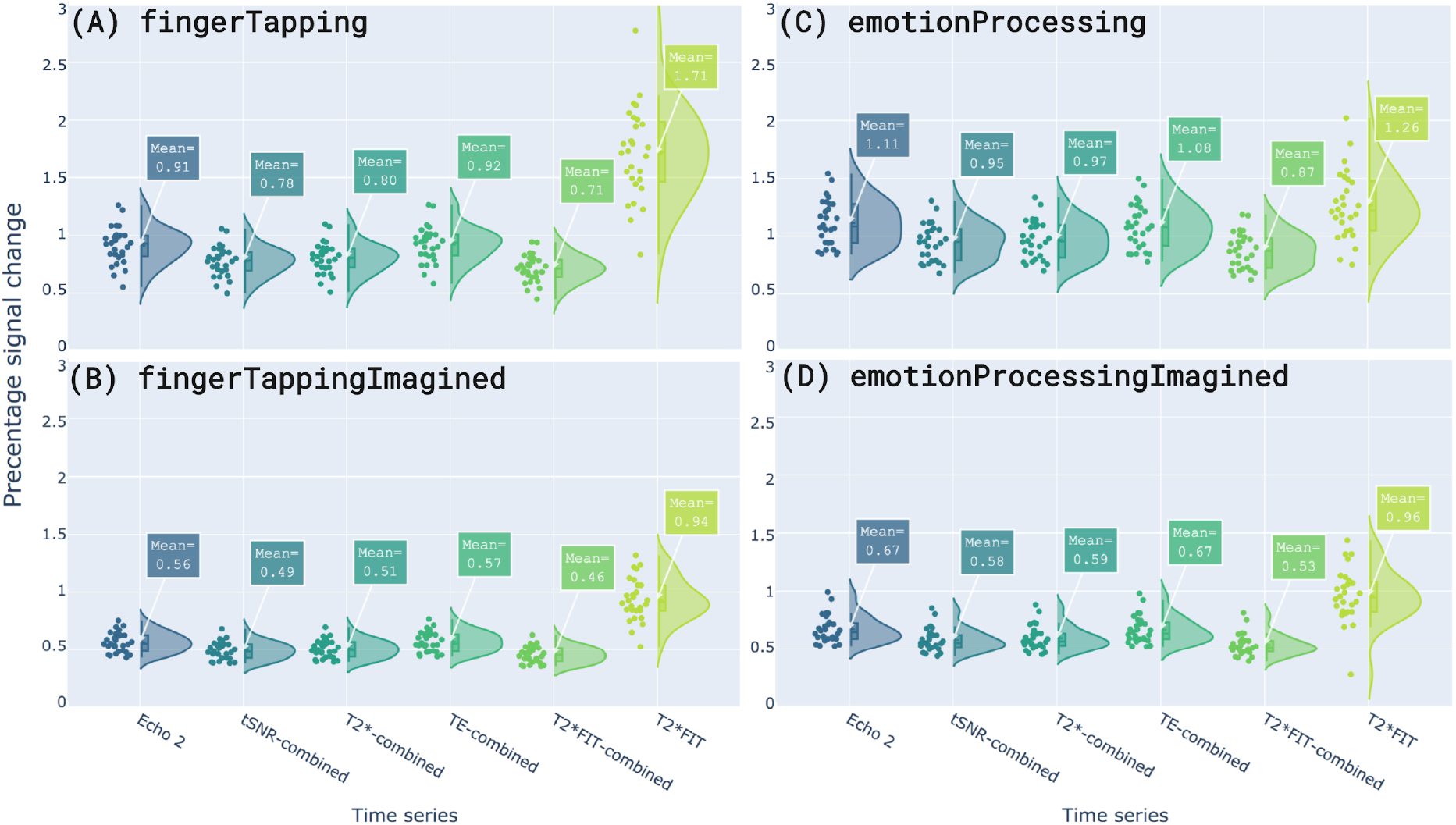
Distribution plots of mean percentage signal change (PSC) values in the FWE-OR cluster for each of the four task runs. Plots from top to bottom are: (A) fingerTapping, (B) fingerTappingImagined, (C) emotionProcessing, and (D) emotionProcessingImagined. PSC values are shown for all six time series, from left to right: Echo 2, tSNR-weighted combination, T2*-weighted combination, TE-weighted combination, T2*FIT-weighted combination, and T2*FIT. For all tasks, the T2*FIT time series effect sizes show mean increases above the effect sizes of the Echo 2 time series, while all multi-echo combined time series effect sizes show similar or decreased means.

Accompanying the above PSC values, Fig. 8 shows distribution plots (over all subjects) of the mean T-statistic values in the respective **FWE-OR** clusters for all task runs: *fingerTapping* (Fig. 8A), *fingerTappingImagined* (Fig. 8B), *emotionProcessing* (Fig. 8C), and *emotionProcessingImagined* (Fig. 8D). For all tasks, it is evident that resulting T-values for the combined echo time series are very similar in size and distribution to that of the Echo 2 time series, while T-values for the *T*_*2*_*\*FIT* time series are notably lower. The low mean T-values of *T*_*2*_*\*FIT* are due to the noise captured when estimating *T*_*2*_*\** per-volume using only three data points, where noisy data would increase standard deviation and decrease the resulting T-values. This is substantiated by the large decrease in tSNR we saw for the *T*_*2*_*\*FIT* time series compared to that of the Echo 2 time series in Figs. 4, 5 and 6. Additionally, the TE-combined time series show slightly higher T-values for all tasks compared to other combined time series. However, this slight increase does not persist when analysing other clusters (e.g. **FWE** or **atlas-based**) as can be viewed with the supplementary browser-based application.

**Figure 8:**
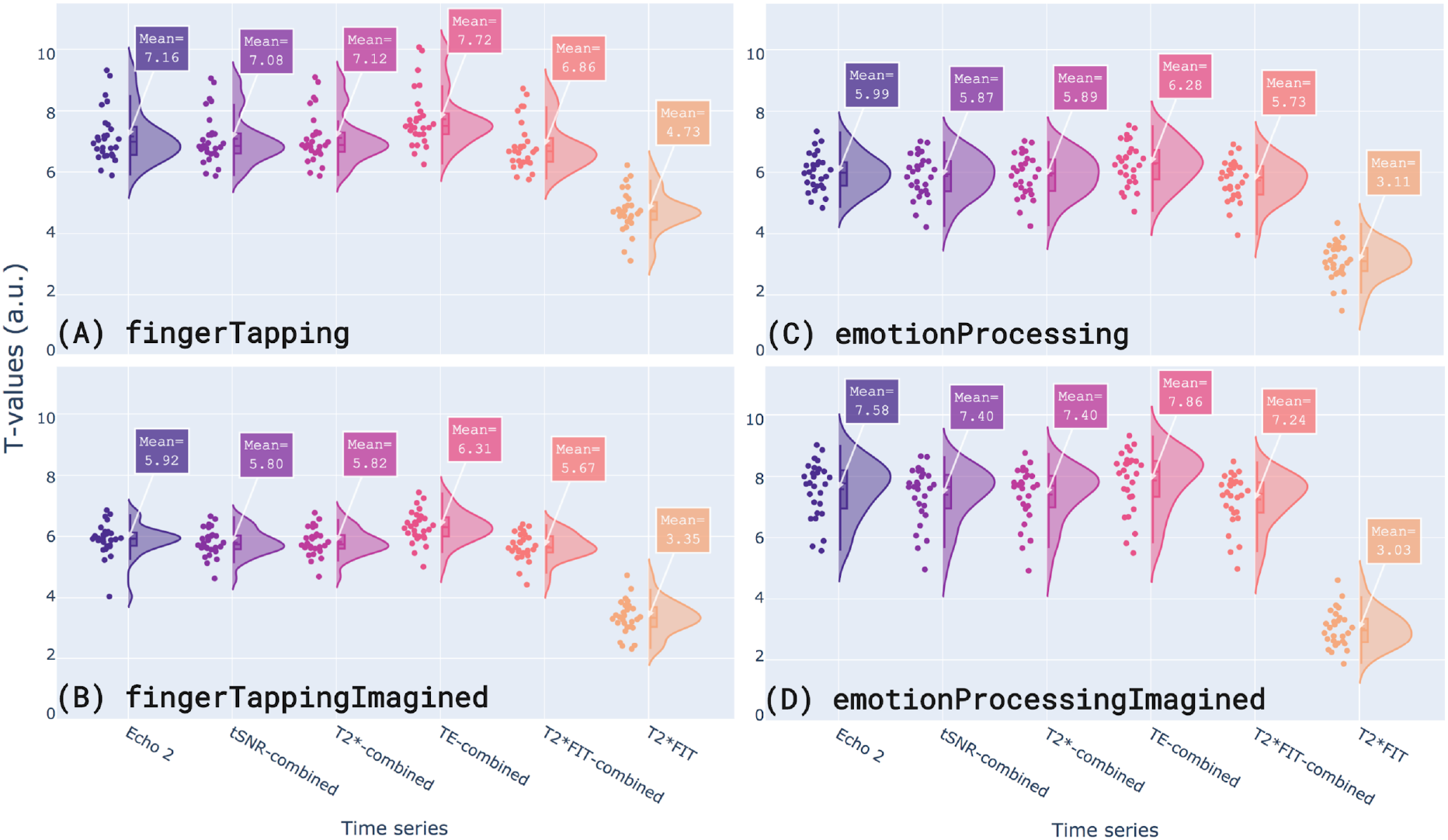
Distribution plots of mean statistical T-values in the FWE-OR cluster for each of the four task runs. Plots from top to bottom are: (A) fingerTapping, (B) fingerTappingImagined, (C) emotionProcessing, and (D) emotionProcessingImagined. T-values are shown for all six time series, from left to right: Echo 2, tSNR-weighted combination, T2*-weighted combination, TE-weighted combination, T2*FIT-weighted combination, and T2*FIT. For all tasks, T-values of the combined echo time series are very similar in size and distribution shape to that of the Echo 2 time series, while T-values for the T2*FIT time series are notably lower.

### 4.5. Temporal percentage signal change and functional contrast

Temporal percentage signal change is useful to inspect the per-volume fluctuations of signal in task-related regions, both for offline and real-time scenarios. tPSC in the offline scenario is calculated per volume from minimally processed data, yielding a per-voxel tPSC time series that can be depicted in a carpet plot or used for ROI analysis. tPSC for realtime scenarios is calculated from real-time minimally denoised *ROI-averaged signal* (with regards to the mean of the preceding baseline “OFF” block or with regards to the cumulative total or baseline mean) yielding the real-time ROI-signal typically used in region-based neurofeedback.

#### Offline temporal percentage signal change

Fig. 9 (A through C) shows per-voxel tPSC as **atlas-based** carpet plots for three time series of the *fingerTapping* task in a single subject (*sub-001_task-fingerTapping*): (A) Echo 2, (B) *T*_*2*_*\**-weighted combination, and (C) *T*_*2*_*\*FIT. T*_*2*_*\**-weighted combination was included as a representative combined time series, since all multi-echo combined time series showed similar intensity fluctuations. Visual inspection indicates that the *T*_*2*_*\*FIT* time series has larger intensity differences in the ON/OFF bands than other signals, suggesting increased functional contrast for *T*_*2*_*\*FIT* compared to single-echo and combined multi-echo time series. This is reflected further in Fig. 9D, which shows averaged tPSC signals within the same atlas-based ROI.

**Figure 9:**
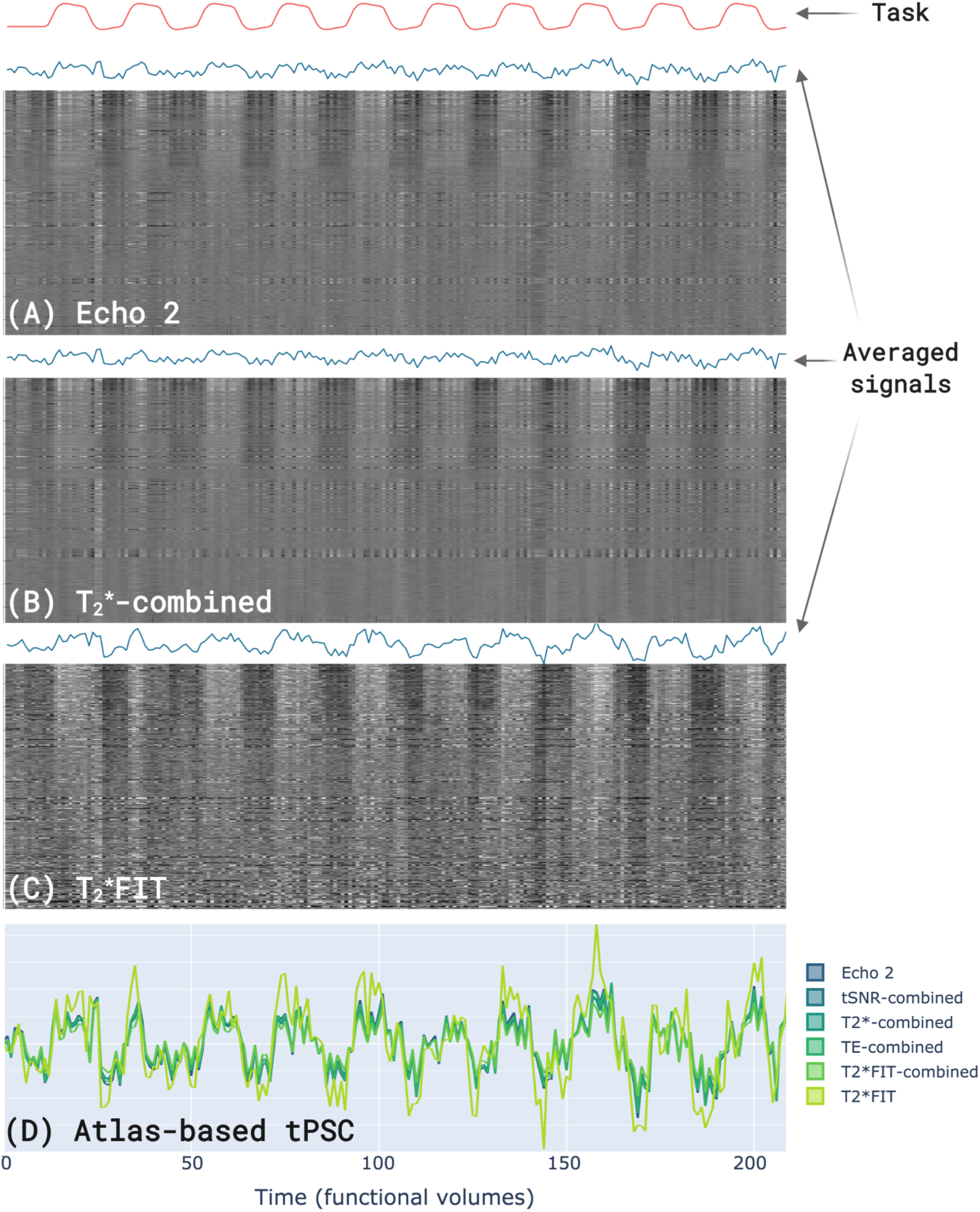
Carpet plots of voxels (A,B,C), and temporal percentage signal change plots (D), in the Atlas-based region (left motor cortex) for the fingerTapping task of a single participant. Carpet plots are shown for the three time series: (A) Echo 2, (B) T2*-weighted combination, and (C) T2*FIT. The increased functional contrast for the T2*FIT time series is clearly visible in the increased light-dark contrast in Fig. 9C. In Fig. 9D, region-based time series signals of temporal percentage signal change are colour coded for Echo 2, tSNR-weighted combination, T2*-weighted combination, TE-weighted combination, T2*FIT-weighted combination, and T2*FIT. Signals are shown as calculated from within regions: (A) FWE, (B) FWE-OR, and (C) Atlas-based. Increased functional contrast for the T2*FIT time series is again visible as higher tPSC amplitudes during task blocks and lower amplitudes during resting blocks.

The *T*_*2*_*\*FIT* signal clearly has a larger functional contrast (higher tPSC during task blocks and lower tPSC during resting blocks) than all other signals, for which the amplitude differences are near-identical. Functional contrast in tPSC signals increase as the regions change from **atlas-based**, to **FWE-OR**, to **FWE** (i.e. as regions become more spatially matched to the participant’s functional activation localisation), as can be seen in the supplementary browser-based application. Note that the *T*_*2*_*\*FIT* time series exhibits a noisier tPSC signal, as can be seen from the larger volume-to-volume fluctuations compared to the overall signal amplitude, visible in both Figs. 9C and 9D.

However, there is a disparity between regions and activity types when assessing the tendency of *T*_*2*_*\*FIT* to increase functional contrast in offline tPSC. Specifically in the amygdala, influenced by increased physiological noise and signal dropout, we are less likely to find functional runs where the *T*_*2*_*\*FIT* time series yields tPSC values with larger functional contrast than the other time series. To demonstrate this quantitatively for all participants, the functional contrasts of the tPSC signals are shown in Fig 10 for the *fingerTapping* and *emotionProcessing* tasks, for the **FWE-OR** and **atlas-based** clusters. For example, for *fingerTapping* in the **FWE-OR** cluster, the percentage increase in functional contrast of *T*_*2*_*\*FIT* time series versus Echo 2 is 100% (from 0.42 to 0.84), whereas the corresponding increase for *emotionProcessing* is 42.86% (from 0.21 to 0.30). Additionally, for *emotionProcessing* in the **atlas-based** cluster, the *T*_*2*_*\*FIT* time series yields a substantial increase in negative task contrasts.

**Fig. 10:**
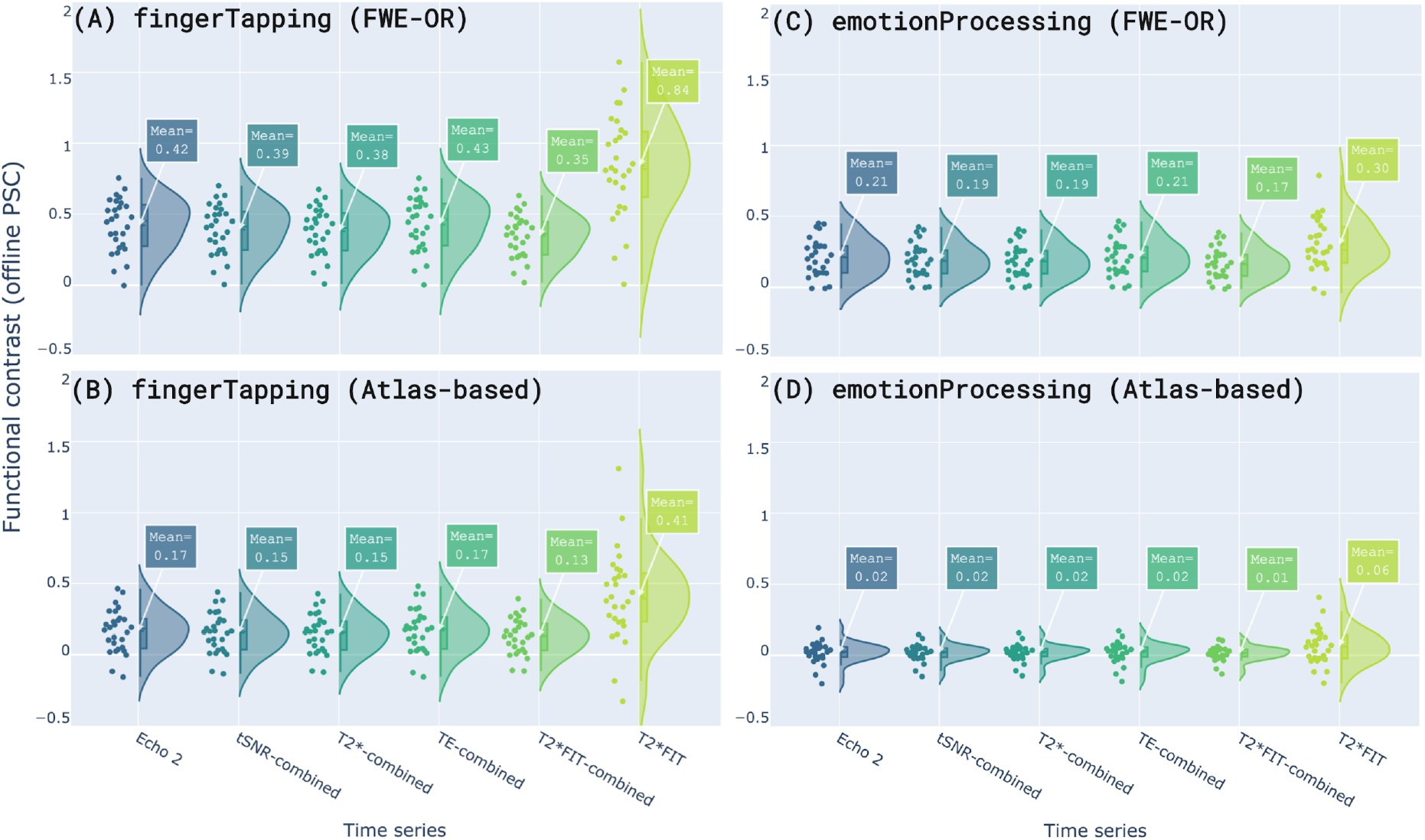
Distributions of functional contrasts calculated from offline temporal percentage signal change of the fingerTapping and emotionProcessing tasks. Contrast distributions are shown for both tasks within the FWE-OR cluster (A and C) and within the atlas-based cluster (B and D). Signals are colour coded for Echo 2, tSNR-weighted combination, T2*-weighted combination, TE-weighted combination, T2*FIT-weighted combination, and T2*FIT. For both tasks, the functional contrast for the T2*FIT time series is greater than the contrasts for all other time series, in both clusters. However, this difference is less pronounced for the emotionProcessing task than for the fingerTapping task. Functional contrast is presented as differences in percentage signal change (y-axes).

#### Real-time temporal percentage signal change

While offline tPSC is useful for post-hoc inspection of signal quality and task activity, it does not accurately reflect the effects seen for real-time scenarios where per-volume calculations can only use information available up to the most recently acquired volume. For that purpose, minimally processed data are cumulatively detrended and real-time tPSC is then calculated with regards to a cumulative baseline mean, which also allows closer comparison with offline tPSC which underwent similar steps. Fig 11 shows functional contrast calculated from real-time tPSC signals for the same tasks and clusters as in Fig. 10.

**Fig. 11:**
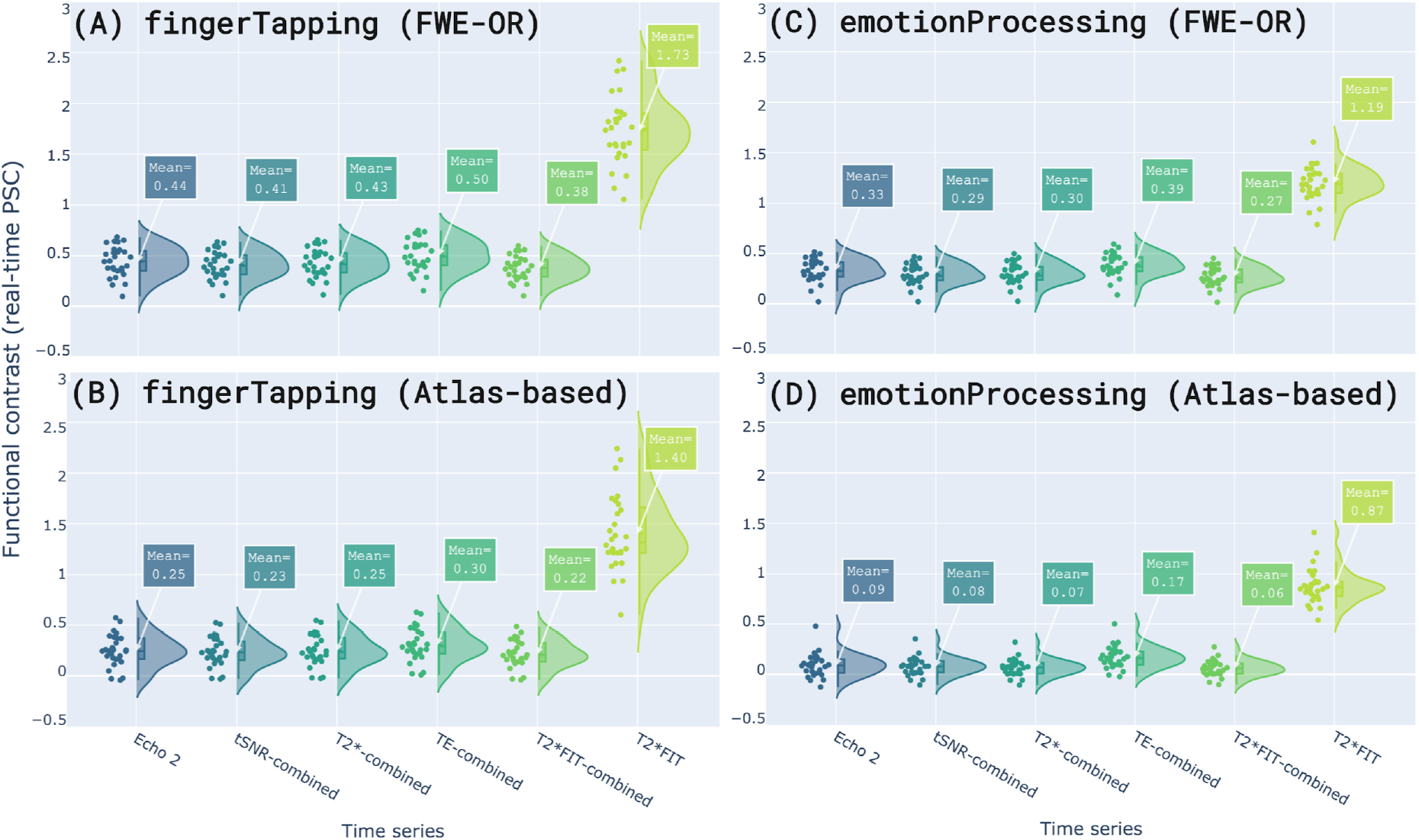
Distributions of functional contrasts calculated from real-time temporal percentage signal change of the fingerTapping and emotionProcessing tasks. Contrast distributions are shown for both tasks within the FWE-OR cluster (A and C) and within the atlas-based cluster (B and D). Signals are colour coded for Echo 2, tSNR-weighted combination, T2*-weighted combination, TE-weighted combination, T2*FIT-weighted combination, and T2*FIT. Similar to the offline tPSC case, the functional contrast (in both tasks) for the T2*FIT time series is greater than the contrasts for all other time series, in both clusters, although this is less pronounced for the emotionProcessing task than for the fingerTapping task. Notably, functional contrast for the real-time T2*FIT time series is substantially increased compared to its offline counterpart (Fig. 10). Functional contrast is presented as differences in percentage signal change (y-axes).

Note the increased functional contrast for the real-time *T*_*2*_*\*FIT* time series (Fig. 11) compared to its offline counterpart (Fig. 10). This is evident for all time series and clusters shown in Fig. 11, where the minimum percentage increase of *T*_*2*_*\*FIT* functional contrast over Echo 2 functional contrast is 260.61% (from 0.33 to 1.19) in the **FWE-OR** cluster of the *emotionProcessing* task. A caveat here, as with the offline tPSC and functional contrast calculations, is that the *T*_*2*_*\*FIT* time series has large overall fluctuations compared to the other time series. It is conceivable that these fluctuations could drive an increase in the estimated functional contrast, regardless of the variability of such an estimate. To take this into account, the functional contrasts are divided by the standard deviation of the tPSC time series to yield the temporal contrast-to-noise ratio (tCNR). This is shown for the **FWE-OR** clusters in the *fingerTapping* and *emotionProcessing* tasks in Fig. 12 below, along with examples of single-participant real-time tPSC signals for the same tasks and clusters. These plots highlight both functional contrast and volume-to-volume fluctuations.

**Fig. 12:**
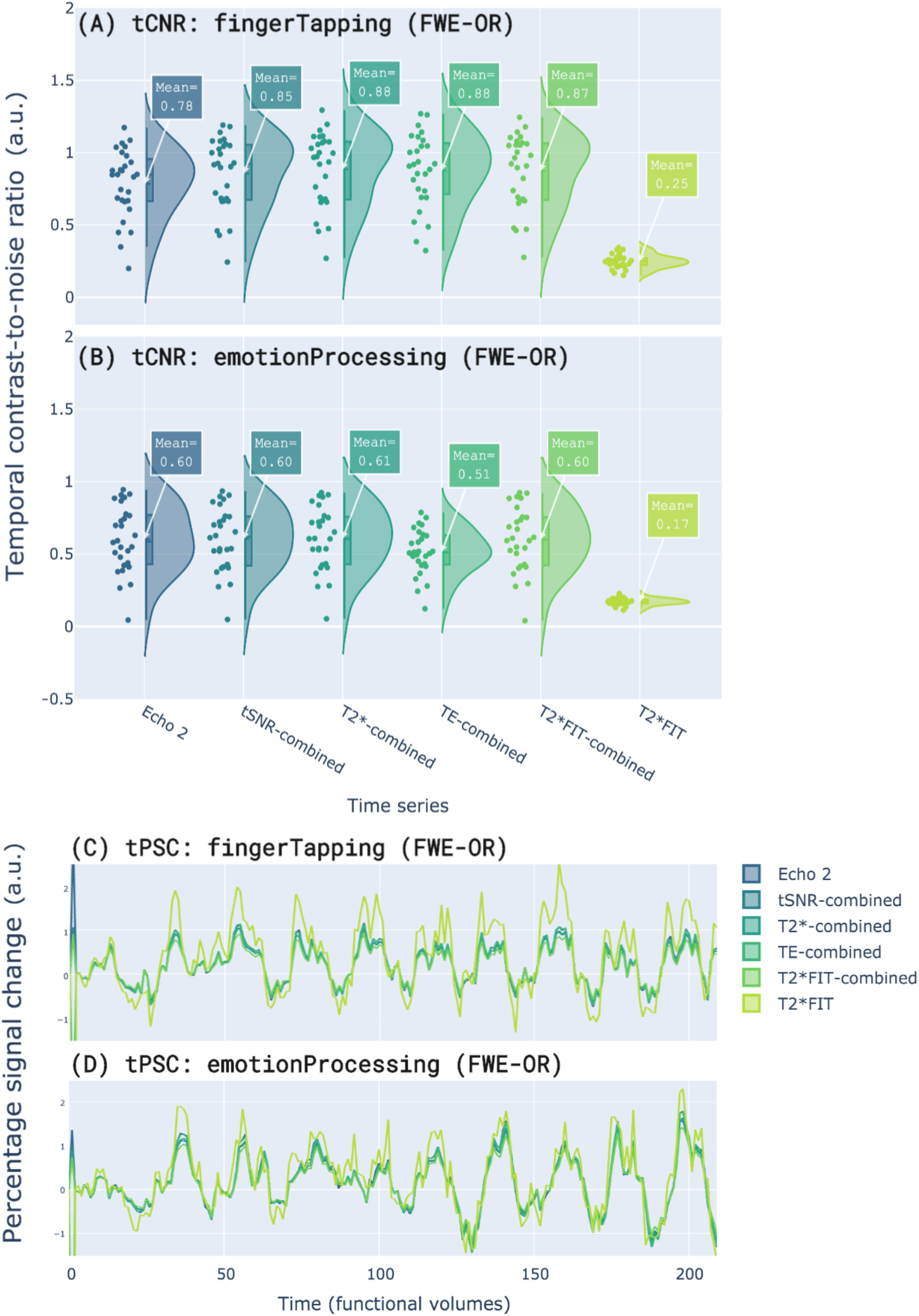
Distributions of mean functional contrast-to-noise ratio (calculated from real-time temporal percentage signal change) of the fingerTapping and emotionProcessing tasks within the FWE-OR cluster. tCNR distributions are shown for the (A) fingerTapping task and (B) emotionProcessing task. Signals are colour coded for Echo 2, tSNR-weighted combination, T2*-weighted combination, TE-weighted combination, T2*FIT-weighted combination, and T2*FIT.

The tCNR distributions clearly show the effect that high signal fluctuations (i.e. increased time series standard deviation) can have on the tCNR. In both Fig. 12A and Fig. 12B, the tCNR of the real-time *T*_*2*_*\*FIT* time series is substantially lower than that of Echo 2 (decreasing by about 70% in both cases) as well as that of all other time series. In Fig. 12C and 12D, tPSC signals in the FWE-OR clusters show higher amplitude differences for the *T*_*2*_*\*FIT* time series compared to all other time series, echoing the increased functional contrast seen for the group in Fig. 11, although an increase in volume-to-volume fluctuations relative to the signal amplitude is also evident. This pattern of increasing *T*_*2*_*\*FIT* time series functional contrast and decreasing tCNR repeats for all tasks and clusters, which shows the importance of taking into account both signal fluctuations and amplitude differences.

## 5. Discussion

In this work we presented a comprehensive exploration and evaluation of existing and novel multi-echo combination and *T*_*2*_*\**-mapping methods for both real-time and offline BOLD sensitivity improvements. A resting state and task-based healthy participant dataset was collected, curated and made available to the community for future investigations. In this dataset, we investigated five time series derived from multi-echo data and their differences from a single echo time series (Echo 2): *T*_*2*_*\**-weighted combination, tSNR-weighted combination, TE-weighted combination, *T*_*2*_*\*FIT*-weighted combination, and the *T*_*2*_*\*FIT* time series. These differences were explored in terms of: temporal signal-to-noise ratio, percentage signal change as task-based effect size measure, offline and real-time temporal percentage signal change, functional contrast, and temporal contrast-to-noise ratio.

Our results, across 28 participants, are summarised as follows. Dropout recovery is more pronounced (in orbitofrontal, ventromedial prefrontal regions as well as inferior and anterior temporal regions) for the *T*_*2*_*\**-weighted, tSNR-weighted, and *T*_*2*_*\*FIT*-weighted combinations than for the TE-weighted combination. All multi-echo combined time series yield increases in tSNR compared to Echo 2, with the newly-proposed *T*_*2*_*\*FIT*-weighted combination resulting in the largest increase in mean tSNR. For the *T*_*2*_*\*FIT*-weighted combination, increases in mean tSNR are larger for the amygdala than for the left motor cortex or the whole brain. In contrast, the *T*_*2*_*\*FIT* time series results in a substantial mean decrease in tSNR from Echo 2. Alternatively, the *T*_*2*_*\*FIT* time series yields the largest effect size measures across all investigated functional tasks and clusters, whereas the effect size measures derived from combined echo time series tend to decrease slightly from those of Echo 2, for all functional tasks. Based on temporal percentage signal change calculated offline from minimally processed data, the *T*_*2*_*\*FIT* time series yields the highest functional contrast for all tasks. Similarly, based on temporal percentage signal change calculated in simulated real-time from cumulatively denoised data, the *T*_*2*_*\*FIT* time series also yields the highest functional contrast for all tasks, although this increase is substantially more than the increase seen for its offline counterpart. For both offline and real-time scenarios, the temporal contrast-to-noise ratio of the *T*_*2*_*\*FIT* time series shows a substantial decrease from Echo 2, while that of all other time series are comparable to Echo 2.

The fact that multi-echo combined time series yields increased tSNR compared to single echo data has been widely demonstrated in previous research, and has been repeated here for all combined time series with respect to Echo 2. Additionally, we show that the novel *T*_*2*_*\*FIT*-weighted combination yields the largest increase, replicating our previous results from a different dataset (Heunis et al., 2019). In the amygdala, a mean increase in tSNR of 53.35% was calculated across participants, while the mean increases for the left motor cortex and the whole brain were respectively 31.63% and 36.95%. These differences suggest that multi-echo combination, and in particular *T*_*2*_*\*FIT*-weighted combination, could prove more useful in terms of tSNR for areas traditionally suffering from suboptimal BOLD sensitivity due to their lower local baseline *T*_*2*_*\**-values. On the other hand, improving tSNR in individual regions could also benefit whole-brain methods where spatially distributed ROIs or networks are used as the neurofeedback substrate (e.g. connectivity-based neurofeedback employed by Megumi et al., 2013, or default mode network-based neurofeedback employed by MacDonald, et al., 2017), since this would decrease spatial variability in BOLD effects and could lead to more accurate brain-wide estimates of interest. Note that we did not explore the approach of averaging the echoes (i.e. simple summation) as for instance originally proposed in Posse et al. (2001), but this approach has proven reduced BOLD sensitivity than the rest of combination approaches investigated here.

While not novel, an important aspect demonstrated here was the decrease in tSNR for the *T*_*2*_*\*FIT* time series. Importantly, the fitting procedure used to estimate per-volume *T*_*2*_*\**- and *S*_*0*_-values (assuming a mono-exponential decay curve) yields noisy results that influence the amplitude of the signal fluctuations with respect to the mean, thus increasing the standard deviation and decreasing tSNR. The pitfalls of assuming mono-exponential (as opposed to multi-compartment) decay and using a fitting procedure with few data points (3 in this case) have been described before (Whittall et al., 1999) and remain applicable here. Future work should aim to exploit technical advances such as simultaneous multi-slice imaging to increase the number of echoes acquired per volume, while investigating more robust models of *T*_*2*_*\**-decay.

While tSNR is a useful quantifier of relative spatial signal increases and dropout recovery, it does not paint the complete picture of influences on BOLD sensitivity. To investigate how multi-echo derived data could improve our ability to link BOLD changes to neuronal effects, we employed statistical task-analysis and effect size measures to show the benefits of rapid *T*_*2*_*\**-mapping over single echo fMRI. For all tasks, the *T*_*2*_*\*FIT* time series consistently yielded the largest standardised effect size measures in terms of percentage signal change calculated offline from contrast maps after participant-level GLM analysis, while the effect sizes for multi-echo combined data decreased slightly. This phenomenon of decreased effect sizes has been reported before for both optimally combined as well as MEICA-denoised data by Gonzalez-Castillo et al. (2016). This was reported for 5 subjects performing an auditory task in a 20 s ON/OFF block paradigm similar to the one in this work. Gonzalez-Castillo et al. calculated per-volume *T*_*2*_*\**-maps (i.e. *T*_*2*_*\*FIT* time series) using the same log-linear fitting approach but with only two echoes (TE = 31.7 ms and 49.5 ms), also in accordance with Beissner et al. (2010), and found that the activation extent, effect sizes and T-statistic values all decreased for the *T*_*2*_*\** time series compared to the original single echo time series. In contrast, we observe that the effect sizes calculated from the *T*_*2*_*\*FIT* time series *increase*, while the related T-statistic values decrease. This difference in the change of the effect size with respect to Echo 2 might be explained by the use of three echoes in our calculation of the *T*_*2*_*\*FIT* time series, instead of two echoes, that could result in reduced accuracy of the *T*_*2*_*\** estimates.

The *T*_*2*_*\*FIT* time series also consistently yielded the largest functional contrasts in terms of differences in task vs. baseline amplitudes in tPSC signals calculated from offline and real-time data. As an example, we observe a 87.91% increase in mean PSC (*T*_*2*_*\*FIT* compared to Echo 02) for the **FWE-OR** cluster of the *fingerTapping* task, and increases in functional contrast for the same task and cluster of 100% and 293%, respectively for offline and real-time scenarios. Interestingly, functional contrast for the real-time calculated tPSC signal showed an increase above the functional contrast calculated from offline data. The main mathematical difference in real-time vs offline approaches that this could be ascribed to is the cumulative calculation vs offline calculation, especially as regards the mean (cumulative baseline mean vs full time series mean).

Theoretically, we should expect an increase in BOLD sensitivity when analysing quantified *T*_*2*_*\** fluctuations versus fluctuations in single echo image intensity, since the separation of *T*_*2*_*\**- and *S*_*0*_ should remove (to a considerable extent) system-level, inflow, and subject-motion effects from the *T*_*2*_*\**-signal. What is left in the form of voxel-based *T*_*2*_*\*FIT*-values would then theoretically be more indicative of local neuronal activity than information derived from single echo data, assuming noise from the fitting procedure and other confounding factors do not attenuate this contrast substantially. Having said that, this highlights an important caveat related to the interpretation of increased functional contrast of the *T*_*2*_*\*FIT* signal compared to that of other signals: the size of signal fluctuations should be taken into account. This is evidenced by the large decrease in temporal contrast-to-noise ratio of the offline and real-time calculated tPSC signals. Temporal smoothing or statistical filtering (e.g. a Kalman filter) are methods that have been employed in neurofeedback tools to smooth out ROI signals calculated in real-time, and this could be investigated as a way to diminish detrimental effects of *T*_*2*_*\*FIT* fluctuations.

In terms of practical applicability to real-time fMRI research, we have shown the usefulness of multi-echo for real-time use cases in a 28-person dataset with several functional task designs. We demonstrate that real-time *T*_*2*_*\*FIT*-weighted combination yields brain wide mean tSNR increases and improves signal recovery in regions affected by dropout, compared to single echo and other combined multi-echo time series. We show additionally that the real-time *T*_*2*_*\*FIT* time series yields large functional contrasts compared to single echo or combined multi-echo time series. These improvements could benefit both real-time brain wide connectivity measures and real-time region-based signals, respectively, showing the possible utility for studies on adaptive paradigms and neurofeedback.

It remains important to consider caveats before implementation and in order to drive future improvements. We noted, as did Clare et al. (2001), that the selection of the region of interest within which to investigate activation effects, functional contrast, tSNR and more, can increase the variability of results and subsequent inferences. This issue was evaluated here considering three different ways of delineating the region of interest: **FWE, FWE-OR**, and **atlas-based**, and we observed attenuation of effect sizes, T-values and functional contrast as regions become less spatially matched to participants’ functional activation localisation. This is particularly important for the real-time neurofeedback context, where a predefined subject-specific region of interest is often required to enable real-time region-based signal extraction.

This concern about variability in the performance due to ROI definition extends to the implementation of real-time denoising steps as well, as noted in our previous work on denoising steps in neurofeedback studies (Heunis et al., 2020b). In the current study, we intentionally implemented a minimal real-time processing pipeline to avoid confounding the results. While the presented benefits of multi-echo fMRI for real-time experiments are promising, further work is necessary to quantify the effects of a full multi-echo and denoising pipeline on BOLD sensitivity and data quality. Taking into consideration the caveats discussed here, we advise researchers planning real-time fMRI studies to design and conduct effective pilot studies and to evaluate the effects robustly before deciding on the optimal multi-echo implementation settings.

Lastly, we have shown that real-time multi-echo processing, specifically rapid *T*_*2*_*\**-mapping and subsequent multi-echo combination is technically viable and practically supported. The software tools generated through this work (and shared with the community) support several per-volume or real-time multi-echo processing operations, including real-time 3D realignment of multi-echo data, real-time estimation of multi-echo decay parameters, real-time multi-echo combination using several weighting schemes, and multiple standard real-time preprocessing steps. It provides a practical toolkit for exploring real-time multi-echo fMRI data and for comparing the effects of acquisition and processing settings on BOLD sensitivity in individuals. Additionally, the interactive browser application allows easy access to the results (https://rt-me-fmri.herokuapp.com/), while the provision of supporting material and code (https://github.com/jsheunis/rt-me-fMRI) allows the presented results to be reproduced and allows replication attempts to be conducted on future datasets.

## 6. Competing Interests

RL, WH, and MB are, respectively, employees of Philips Research and Philips Healthcare in The Netherlands. The other authors have declared that no further competing interests exist.

## 7. Acknowledgements

This work was funded by the foundation Health-Holland LSH-TKI (grant LSHM16053-SGF) and supported by Philips Research. LH was supported by the European Union’s Horizon 2020 research and innovation program under the Grant Agreement No 794395. CCG was supported by the Spanish Ministry of Economy and Competitiveness (Ramon y Cajal Fellowship, RYC-2017-21845), the Basque Government (PIBA_2019_104) and the Spanish Ministry of Science, Innovation and Universities (MICINN; PID2019-105520GB-100).

## Notes

https://dataverse.nl/dataverse/rt-me-fmri

https://rt-me-fmri.herokuapp.com

https://github.com/jsheunis/rt-me-fMRI

